# Design, implementation, and functional validation of a new generation of microneedle 3D high-density CMOS multi-electrode array for brain tissue and spheroids

**DOI:** 10.1101/2022.08.11.503595

**Authors:** Lisa Mapelli, Olivier Dubochet, Mariateresa Tedesco, Giacomo Sciacca, Alessandra Ottaviani, Anita Monteverdi, Chiara Battaglia, Simona Tritto, Francis Cardot, Patrick Surbled, Jan Schildknecht, Mauro Gandolfo, Kilian Imfeld, Chiara Cervetto, Manuela Marcoli, Egidio D’Angelo, Alessandro Maccione

## Abstract

In the last decades, planar multi-electrode arrays (MEAs) have been widely used to record activity from *in vitro* neuronal cell cultures and tissue slices. Though successful, this technique bears some limitations, particularly relevant when applied to three-dimensional (3D) tissue, such as brain slices, spheroids or organoids. For example, planar MEAs signals are informative on just one side of a 3D-organized structure. This limits the interpretation of the results in terms of network functions in a complex structured and hyperconnected brain tissue. Moreover, the side in contact with the MEAs often shows lower oxygenation rates and related vitality issues. To overcome these problems, we empowered a CMOS high-density multi-electrode array (HD-MEA) with thousands of microneedles (μneedles) of 65-90 μm height, able to penetrate and record in-tissue signals, providing for the first time a 3D HD-MEA chip. We propose a CMOS-compatible fabrication process to produce arrays of μneedles of different widths mounted on large pedestals to create microchannels underneath the tissue. By using cerebellar and cortico-hippocampal slices as a model, we show that the μneedles efficiently penetrate the 3D tissue while the microchannels allow the flowing of maintenance solutions to increase tissue vitality in the recording sites. These improvements are reflected by the increase in electrodes sensing capabilities, the number of sampled neuronal units (compared to matched planar technology), and the efficiency of compound effects. Importantly, each electrode can also be used to stimulate the tissue with optimal efficiency due to the 3D structure. Furthermore, we demonstrate how the 3D HD-MEA can efficiently penetrate and get outstanding signals from *in vitro* 3D cellular models as brain spheroids. In conclusion, we describe a new recording device characterized by the highest spatio-temporal resolution reported for a 3D MEA and significant improvements in the quality of recordings, with a high signal-to-noise ratio and improved tissue vitality. The applications of this game-changing technique are countless, opening unprecedented possibilities in the neuroscience field and beyond.

## Introduction

Planar multi-electrode arrays (MEAs) are widely used to perform multisite extracellular recording of brain activity *in vitro* and *ex vivo*. Other than on primary or IPSC-derived 2D cultures (Amin et al., 2016, 2017; Cutarelli et al., 2021; Sun et al., 2022) these devices are also employed on structured 3D models as acute brain slices and explanted tissues (Mapelli and D’Angelo, 2007; González-Calvo et al., 2021; Gagliano et al., 2022; Hu et al., 2022; Mahadevan et al., 2022; Tognolina et al., 2022), and 3D culturing techniques as spheroid or organoids (Georgiou et al., 2020; Mayer et al., 2020; Dorgau et al., 2022). However, using planar MEAs with 3D tissue bears some critical limitations. The recorded signal derives from the outermost layers, with little or no access to the activity of the cells inside the 3D structure. This is even more relevant in acute slices (Buskila et al., 2015) for the presence of a dead cells layer caused by the cutting procedure, and in organoids (Tasnim and Liu, 2022) where a scaffold matrix (e.g., Matrigel) often encapsulates the tissue (additionally providing insulation in many cases). Several approaches have been developed in the MEA field in the effort to produce penetrating electrodes capable of retrieving signals from the bulk of the 3D model, thus overcoming the main issues. First attempts to manufacture 3D electrodes have been already presented in the 90s: Hoogerwerf et al. (Hoogerwerf and Wise, 1994) implemented an 8 x 16 electrodes grid spaced at 200 μm assembled on a CMOS device. Similar approaches were proposed in (Thiébaud et al., 1997; Bai et al., 2000; Heuschkel et al., 2002; Irons et al., 2008; Du et al., 2009; Musick et al., 2009). In these works, 3D electrodes were manufactured on a glass substrate producing only a few sensing sites with a poor spatial resolution of hundreds of micrometers. In Aziz et al., 2007, standard bonding equipment has been used to fabricate 16 x 16 gold microelectrodes with 170 μm pitch. The studs are manufactured with a base diameter of 80 μm and a height of approximately 100 μm. In this work, electrodes of different heights were produced using standard fabrication techniques, but the need to process the electrodes one after the other makes this approach quite unrealistic for large-scale adoption in research. In,Hashemi et al., 2010, electrodes with a much smaller pitch were realized with a stack of metal 1 to metal 5 patches with all intermediate vias. Then, the SiO_2_ layer was removed down to metal 3, leaving electrodes with a height of only a few micrometers, not enough to efficiently penetrate any tissue. In Gunning et al., 2013, the authors proposed an alternative fabrication method where the sidewalls of tapered holes in Si substrate are covered with insulator and metallic layers and filled with parylene. Si was then etched on the opposite side to expose the tips of the needles. This process was used to produce a 512 grid of electrodes up to 250 μm in height and spaced at 60 μm. The drawback of this approach is the double-side Si wafer process, which makes this microfabrication process extremely complex. Currently, 3D electrodes on standard glass MEAs for *in vitro* and *ex vivo* applications are also commercially available from Multichannel Systems (Multichannel Systems 2018 Microelectrode array (MEA) manual, 2018; Vernekar and Laplaca, 2020). They provide standard passive MEAs with 60 electrodes with a pitch varying from 100 to 250 μm and a height from 40 to 100 μm.

Despite the improved signal quality in the recordings, for both spiking activity and local field potentials (LFP), the currently available devices still provide a large pitch and low-count number of recording sites. These features pose substantial limitations in their use with brain tissues or 3D models, which require recordings from large interconnected areas of several mm^2^, still keeping enough spatial resolution to discriminate signal sources within small anatomical compartments (Egert et al., 2002; Ma et al., 2008). Furthermore, 3D models need efficient nutrient diffusion, proper oxygenation, and fast metabolic waste removal to avoid rapid necrosis of the core of the tissue (Liu et al., 2020). Conventional MEA devices require the tissue to be in intimate contact with the surface containing the electrodes, thus strongly limiting the solution exchange at the recorded layers, often resulting in a rapid decrease of the activity or poorly physiological experimental conditions. To avoid such limitations, perforated MEAs have been proposed, allowing liquid exchange from the bottom of the device (Killian et al., 2016) alternatively, an interesting approach from Rowe et al., 2007 proposed an 8×8 matrix of microstructured SU-8 pillars, each of them providing two electrodes and multiple fluidic ports. Still, in both approaches, the number of recording sites remains low, with poor spatial resolution.

In this paper, we introduce for the first time a new fabrication technique able to provide a chip with thousands of microneedles (μneedles) of height ranging from 60 to 90 μm and width from 14 to 26 μm with 60 μm pitch. At the bottom of each needle, a larger pedestal is structured. This determines the formation of a tight matrix of microchannels below the tissue penetrated by the needles, avoiding direct contact with the basement of the chip. Notably, this fabrication process is fully CMOS-compatible, allowing the generation of the first high-density MEAs based on this technology (Berdondini et al., 2009) The result is a monolithic CMOS high-density MEA providing a 64 x 64 μneedle electrode grid with an integrated microfluidic system to guarantee efficient oxygen, nutrient, and chemical diffusion at the bottom layers of the tissue. Such technology condenses penetrating capability, high-resolution recording, and improved tissue viability in a unique device. The system efficiency was validated by recording spontaneous, chemically modulated, and electrically evoked activity from cerebellar and cortico-hippocampal brain slices. The data obtained with the 3D chips were compared to planar high-density MEAs on the same samples, showing a significant improvement in the efficiency of recordings. In addition, smaller needle sizes have been finally validated to record from small 3D engineered tissue as brain spheroids, which typically provide a lower cellular density than *ex vivo* tissue and have typical diameters of a few hundreds of micrometers.

The potential of such technology to provide groundbreaking access to advanced 3D model systems has direct implications for basic neuroscience research, biomarker discovery, and pre-clinical evaluation of novel neuromodulating or neuroprotective compounds.

## Materials & Methods

### Microneedle and microchannel fabrication

The fabrication process is based on SUEX photosensitive dry film, and it has been developed to be CMOS-compatible for wafers of 150 mm diameter and thickness > 700 μm. The μneedles were produced in two versions, the first with a needle width of 26 μm and height of 90 μm and the second with thinner needles of only 14 μm but with a decreased height down to 65μm. The first was tested for large dense brain tissues, while the second was validated on low density, small size brain spheroids.

Fig. 1A shows a section of one of the chips on the wafer as it is provided by the CMOS foundry, the top surface is covered by a thin insulator layer of silicon nitride with openings in correspondence of the aluminum electrodes that will be contacted with the μneedles. After removing the oxide layer over the aluminum electrodes by using a back etching process, the wafer is covered with an electrochemical seed layer consisting of a first adhesion metal of TiW followed by a conduction layer made up of copper (Fig. 1B). As shown in Fig. 1C, a layer of SUEX photoresist, with a thickness of 25 μm, is spun and patterned in order to create a mold with openings for pedestal electro-plating. The mold thus created is then electroplated with 99% pure biocompatible gold forming the pedestals. The wafer is then planarized by using a diamond turning process to result in the structure of Fig 1D with a final pedestal height of 15 μm. With reference to Fig. 1E, a further photoresist layer of SUEX, with a thickness in the range of 80-100 μm, is spun and patterned in order to create a mold with openings for μneedle electro-plating. The mold thus created is electroplated with gold as shown in Fig. 1E. The deposited gold provides the conductive needles. The resulting structure similarly to the pedestal formation is then planarized by using a diamond turning process reducing the height of 95 μm for large needles and 70 μm for smaller ones (Fig. 1F). The photoresist molds are then stripped, and the electrochemical seed layer is etched to remove electrical shortcuts between the microneedles resulting in the structure of Fig. 1G. The structure is then covered by vapor deposition with an electrical insulation coating consisting of a 2 μm layer of Parylene-C as shown in Fig. 1H. Using a planarization process with a diamond turning, the insulation coating is removed from the top of the microneedles, to finally obtain an electrode with the electrical sensing area located on the top of the needle only (Fig. 1I). The resulting sizes are 90 μm high for large needles of 26 μm width, and 65 μm high for smaller ones of 14 μm width.

**Fig. 1.**
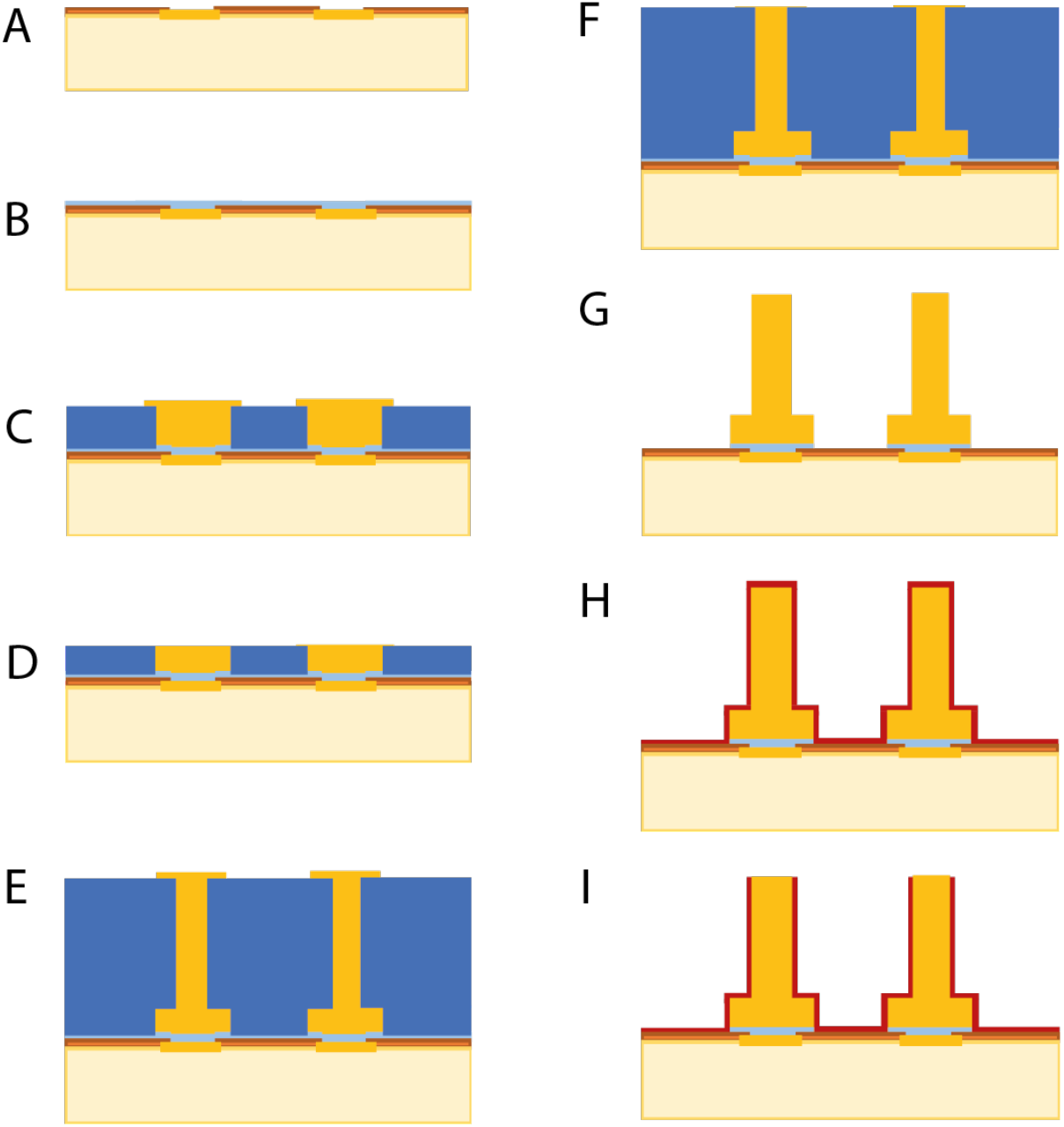
Schematic overview of the fabrication process steps to structure μneedles on top the electrodes of a CMOS HD-MEA. **A)** Section of the HD-MEA showing electrodes (orange) opened in the silicon nitride insulator layer (brown). **B-D)** Lithography steps to structure the large pedestals at the base of the μneedles. **E,F)** Same as B-D) to form the μneedles. **G)** Bare 3D gold μneedle electrodes not electrically insulated. **H,I)** Deposition of an insulating parylene-C layer and opening of the tip of the μneedles.

### Preparation of acute cerebellar and cortico-hippocampal slices

Animal maintenance and experimental procedures were performed according to the international guidelines of the European Union Directive 2010/63/EU on the ethical use of animals and were approved by the local ethical committee of the University of Pavia (Italy) and by the Italian Ministry of Health (following art.1, comma 4 of the D.Lgs. n. 26/2014 approved on December 9th, 2017).

Acute cerebellar and cortico-hippocampal slices were obtained from C57Bl6 mice (20-26 days old, either sex) following a standard procedure, as reported previously (Soda et al., 2019; Tapella et al., 2020; Gagliano et al., 2022). Briefly, mice were deeply anesthetized using halothane (Sigma-Aldrich) and killed by decapitation. Acute 220 μm parasagittal slices of the cerebellar vermis were cut using a vibroslicer (Leica VT1200S, Leica Microsystems). In parallel, acute 320 μm coronal brain slices containing the hippocampus were obtained on a second vibroslicer (Leica VT1200S). During the whole procedure, slices were maintained in ice-cold Krebs solution containing (mM): 120 NaCl, 2 KCl, 1.2 MgSO_4_, 26 NaHCO_3_, 1.2 KH_2_PO_4_, 2 CaCl_2_, and 11 glucose, and was equilibrated with 95% O_2_-5% CO_2_ (pH 7.4). Slices were then recovered for at least 1h in Krebs solution, at room temperature. During recordings, the slices were continuously perfused with Krebs solution (2 ml/min) using a peristaltic pump (Ismatec). When specified, 3 μM TTX (tetrodotoxin, Tocris) was added to the Krebs solution. When specified, for experiments with cortico-hippocampal slices, Krebs solution was modified to increase tissue excitability (increasing [K^+^] to 8 mM, [Ca^2+^] to 4 mM, and abolishing [Mg^2+^]).

### Preparation of brain spheroids

The experimental procedures and animal care complied with the European Communities Parliament and Council Directive of 22 September 2010 (2010/63/EU) and with the Italian D.L. n. 26/2014, and were approved by the Italian Ministry of Health (protocol number 75F11.N.6JI, 08/08/18). All possible efforts were made to minimize animal suffering and the number of animals used.

Rats (Sprague Dawley 200–250 g) were housed at the animal care facility of Department of Pharmacy (DIFAR), University of Genova, Italy, at constant temperature (22 ± 1 °C) and relative humidity (50%), under a light–dark schedule (lights on 7AM–7PM), and with free access to standard pellet diet and water. The development of brain spheroids (neurospheres) from primary embryonic neurons cultures was obtained following standard procedures as reported previously (Dingle et al., 2015, 2020; Khan et al., 2018; Sevetson et al., 2021). Briefly, primary cortexes of rat embryos E18-19 were isolated in HBSS without Ca^2+^ and Mg^2+^ and digested in a solution of 0.125% Trypsin + DNAse 50 μg/ml at 37°C for approximately 18-20′. The proteolytic action of Trypsin was blocked by the addition of 10% FBS medium (Neurobasal + B27 + Glutamax-100 1% + 10 ug/ml Gentamicin). The medium with 10% serum was removed and the tissue washed twice with fresh FBS-free medium. Then, by using a Pasteur pipette with a narrow tip the tissue was mechanically dissociated, counted in a haemocytometer to assess cell yield, and the cell suspension was diluted to the desired concentration. Spheroid structures were obtained by depositing the cells directly into 96-well plates (Biofloat) fabricated with ultra-low attachment plastic. It was possible to generate spheroids of different sizes by increasing the number of cells plated in each well. Typically, 12,000-13,000 cells per well were plated, which at the end of the development period gave rise to spheroids around 500 microns in diameter. Spheroid formation occurs quickly, and 48 hours after plating it was possible to appreciate their presence under the microscope. The plating medium was composed by Neurobasal + B27 + Glutamax-100 1% + 10 μg/ml Gentamicin, after 3 days in culture, 50% of the total volume of medium was replaced (to eliminate any debris) and finally, on day 5 *in vitro*, the medium was replaced 80% with BrainPhys + SM1 + 1%Glutama-100 + 50 μg/ml Gentamicin. The culture of the spheroids was maintained in an incubator at 37 °C, 5% CO2 and 95% humidity for approximately 18-25 days with medium changes at 50% every 2-3 days. Then the individual spheroids were transferred directly onto the 3D high-density multi-electrode array (HD-MEA) chip for acute evaluation of electrical activity.

### Vitality fluorescence analysis

Cerebellar slices were loaded with calcein AM, a live cell-permeant dye, by incubation for 40 min in Krebs solution containing 20 μM calcein AM (acetoxymethyl ester of calcein; ThermoFisher Scientific). The slices were then washed three times in normal Krebs solution and fixed for at least 2 hours at RT or overnight at 4 °C in PBS PAF 4%. Slices were then washed 5 min in PBS for three times, and mounted with Fluoroshield mounting medium with DAPI (Abcam, UK). The 220 μm thick cerebellar slices were mounted between two coverslips in order to analyze both sides; images were then acquired with TCS SP5 II LEICA (Leica Microsystem) equipped with an inverted microscope LEICA DM IRBE (UniPV PASS-BioMed facilities) and analyzed with ImageJ (Schneider et al., 2012). To estimate the percentage of vital granule cells, the ratio of cells loaded with calcein AM vs. granule cells stained only for DAPI (vital + non vital cells) was manually calculated. The cell’s vitality was analyzed on 20 fields per sample, on at least three different planes every two to avoid double counting.

### Voltage-Sensitive Dye imaging (VSDi)

VSDi recordings were performed on acute cerebellar slices, as in (Mapelli, 2010; Gandolfi et al., 2013, 2015; Soda et al., 2019). Briefly, slices for optical recordings were incubated for 30 minutes in oxygenated Krebs solution added to a 3% Di-4-ANEPPS (Invitrogen) stock solution and mixed with an equal volume of fetal bovine serum (Invitrogen), reaching the final dye concentration of 2mM. The slices were then rinsed with Krebs solution and transferred to the mounting stage of an upright epifluorescence microscope (Slicescope, Scientifica Ltd). A proper set of filters (excitation filter *λ* = 535 ± 20nm; dichroic mirror *λ* = 565 nm; absorption filter *λ* > 580 nm) was used to project the light to the slice and acquire the fluorescence signal, passing through a 20X water immersion objective (XLUMPlanFl, 0.95 numerical aperture; Olympus) and collected by a CCD camera (MICAM01 SciMedia, Brain Vision). The imaging system was connected to a PC unit through an input/output interface (Brain Vision), controlling the pattern of illumination, stimulation, and data acquisition. With this asset, the pixel size was 4.5 × 4.5 μm. Fluorescent signals were acquired using BrainVision software, with the sampling rate of 0.5 kHz. The mossy fiber bundle was stimulated with a tungsten bipolar electrode (WPI) connected to a stimulation unit through a stimulus isolator (stimulus intensity 15V, duration 250μs). Single-pulse stimulation was delivered at 0.1Hz and every VSDi trace was obtained from ten repetitions, to improve signal-to-noise ratio (see Soda et al., 2019) Data analysis was performed as previously described (Soda et al., 2019). Firstly, traces were filtered (3 x 3) using both a cubic and a spatial filter. Secondly, ad hoc routines in Matlab (Mathworks) were implemented to analyze the responses to stimulation, as rapid and transient increases in fluorescence exceeding the baseline average *Δ*F/F by more than 2.5 standard deviations. This procedure allowed obtaining maps of the spatial distribution of granular layer responses to stimulation. The percentage of the granular layer activated by the stimulus was calculated dividing the number of pixels showing a response to the stimulation by the total number of pixels sampling the granular layer and multiplying by 100. Statistical comparison was performed using unpaired Student’s *t* test, with significance level <0.05. Data are reported as mean ± SEM (standard error of the mean).

### Electrophysiological recordings

Electrophysiological recordings on cerebellum and cortico-hippocampal brain slices were performed using 2D and 3D high-density multi-electrode arrays (HD-MEA) from 3Brain AG. Both planar electrodes (2D) and structured with μneedles 3D HD-MEAs rely on the same IC CMOS chip. The chip provides 4096 electrodes arranged in a 64×64 grid with an electrode pitch of 60 μm (total recording area 3.8 × 3.8 mm^2^). Each electrode integrates an amplification stage and a multiplexing architecture as presented in Imfeld et al., 2008, enabling simultaneous recordings from all the 4096 electrodes at a sampling rate of 20kHz. Differently from the original architecture, a switching circuitry allows to route each individual electrode to a current signal generator integrated in the acquisition system (BioCAM Duplex from 3Brain AG) enabling electrical stimulation from the HD-MEA chip. Planar electrodes consist of a square of 21 μm side covered by a thin layer of Pt, while the three-dimensional microneedle electrodes (Fig. 2) provide a gold basement 30 μm width and 15 μm height. On top of the pedestals, two versions of the μneedles were realized: a large one of 26 μm width and 90 μm height and a thinner one of 14 μm width and 65 μm height. The electrodes are insulated by a layer of parylene leaving a conductive aperture on the tip (see *Microneedle and microchannel fabrication* section).

**Fig. 2.**
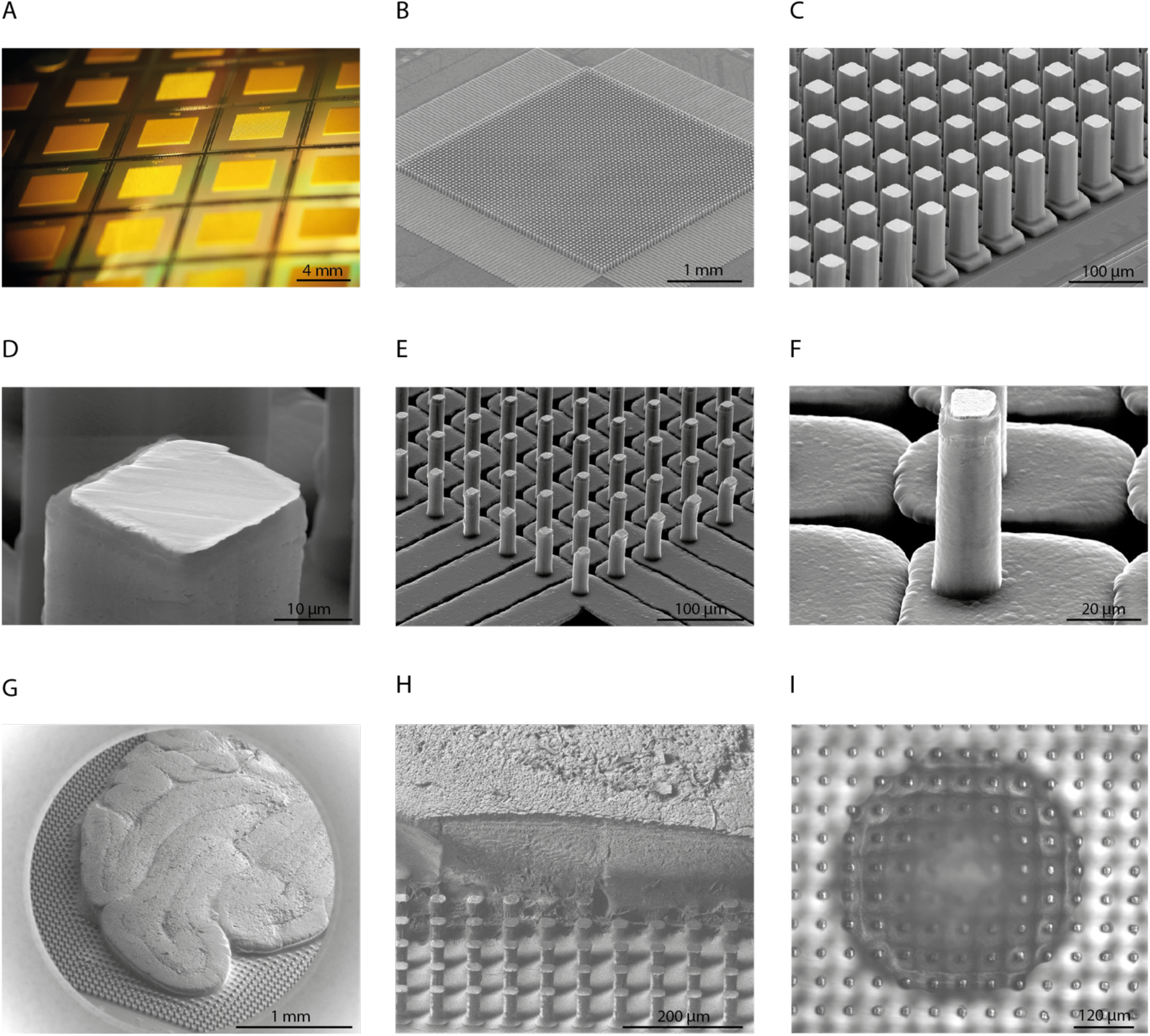
Realization of the 3D μneedle structures integrating at the base a grid of microchannels. **A)** Structuring of the 3D HD-MEA at wafer level before dicing in single chips. **B-H)** Series of Scanning Microscopy Imaging (SEM) showing: **B)** overview of the 4096 μneedle electrodes forming a “bed of nails”, **C,D)** details of the large μneedles compared with the smaller μneedles presented in panels **E,F)**, **G)** a 250μm thick cerebellar slice positioned and fixed on the 3D HD-MEA, **H)** same cerebellar slice as in G) after a longitudinal cut with a Focused Ion Beam showing how the μneedles penetrated the tissue without bending or deforming. **I)** Phase contrast micrograph of a brain spheroid mounted on a 3D HD-MEA showing the intimate attachment of the tissue on the μneedles.

To perform a direct comparison of the efficiency in recording signals of the microneedle electrodes vs. planar ones, each slice was measured on both a 2D and a 3D HD-MEA. A total of 6 cerebellum slices and 4 cortico hippocampal slices were tested from 3 different animals. On the 2D HD-MEA the slice was stabilized using a platinum anchor with wires of 100 μm width, while no holder was used on the 3D HD-MEA. Recordings consisted in 3-5 minutes of spontaneous activity performed 5 minutes after the slice was positioned on the chip.

TTX modulation of spontaneous activity in cerebellar slices was recorded by acquiring activity for about 4 minutes in standard Krebs solution, followed by perfusion of Krebs with 3 μM TTX. Each slice was monitored for the following 3 minutes or until the activity completely disappeared. 10 slices from 3 different animals were tested against TTX for both 2D and 3D HD-MEA.

On some cerebellar slices mounted on a 3D HD-MEA, electrical stimulation was also performed by releasing a biphasic stimulation from two adjacent electrodes selected in the granular layer of a lobule, in correspondence of the mossy fibers. Stimulation intensity was set at 65 μA for 150 μs for the first phase followed, 30 μs after, by a second phase with −50 μA delivered for 40 μs. Different lobules of the same slice were tested.

Spontaneous activity of cortical brain spheroids was measured at DIV 19-20 by using the 3D HD-MEAs. Recordings consisted in acquiring a few minutes of basal activity in acute condition: three spheroids per chip were transferred with a pipette with a large tip on the 3D HD-MEA previously treated with a plasma cleaner process to make the surface hydrophilic. No anchor was used, and the spheroids started showing spontaneous spiking activity immediately after the placement on the chip. Two different dimensions of brain spheroids of about 400 μm and 600 μm were tested. During recording, the spheroids were kept in culture medium with a 5% CO2 humidified flux and at a temperature of approximately 35°C.

### Electrophysiological recordings: data analysis

Data were acquired, visualized, and stored using BrainWave 4 software from 3Brain AG. The software allows superimposing the image of the slice positioned on the chip with the map of activity for anatomically matching the recordings with the brain regions (e.g., Fig. 4A,B). Spikes were extracted from the electrophysiological recordings using the algorithm presented in Muthmann et al., 2015. Detected events were then sorted as described in Hilgen et al., 2017. An active unit was considered for the analysis if the spike rate was higher than 0.5 spike/s in cerebellar slices and 0.1 spike/s in cortico-hippocampal brain slices and if its position was within the considered brain region. For the “*Increased sensing capabilities in detecting spiking activity on brain tissue*” and “*Accelerating compound effect in modulating electrical activity*” data analysis, a total number of 6 and 10 cerebellar slices were respectively considered. All the slices were extracted from 3 different animals. We used 4 cortico-hippocampal brain slices extracted from a single animal. All datasets were initially assessed to ensure the distribution followed a Gaussian curve using the Shapiro–Wilk normality test. The groups comparison of the “*Increased sensing capabilities in detecting spiking activity on brain tissue*” data analysis was performed using an unpaired Student’s *t* test. The “*Accelerating compound effect in modulating electrical activity*” data analysis was conducted using a two-way ANOVA followed by a Tukey’s multiple comparison post-hoc test. Statistical significance was set at *p*<0.05 and all the statistical analyses were performed using GraphPad Prism (9.0, GraphPad Software, La Jolla, CA).

**Fig. 3.**
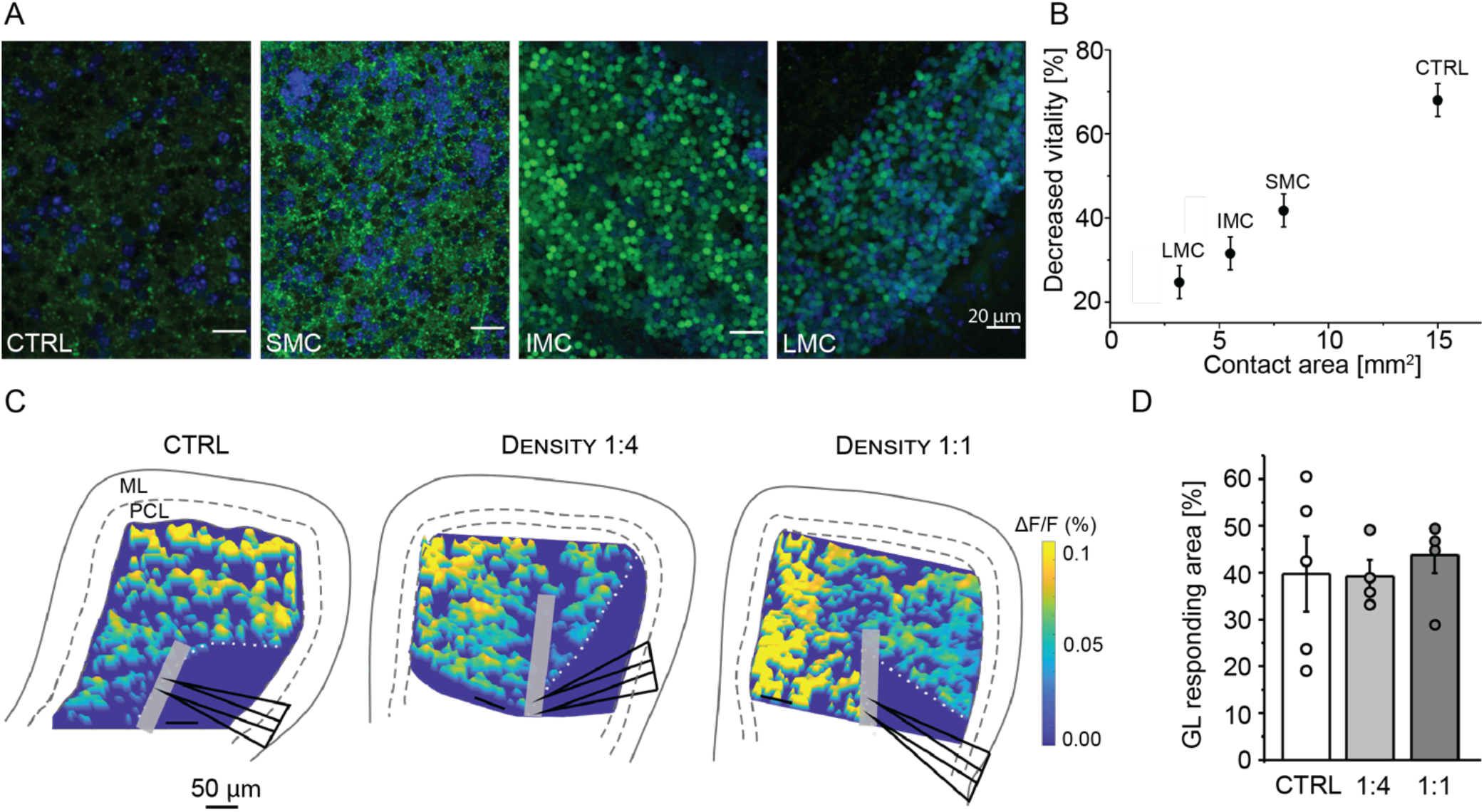
3D HD-MEA chips showed improved tissue vitality without perturbing network functionality. **A)** Representative confocal images at 40x magnification of cerebellar slices loaded with calcein AM, after 1h placement over SMC, IMC, LMC chips, or in control (CT, over the glass chamber). Living cells at the time of calcein exposure are labeled in green; the blue staining identifies cell nuclei (DAPI). Scale bar 20μm. **B)** The plot shows the decreased vitality compared to the contact area for each configuration shown in A). The decreased vitality is calculated by quantifying the calcein stained cells from confocal images on the bottom (on the chip/glass) compared to the top (in contact with Krebs solution) side of the slices. The contact area depends on the microchannel widths in SMC, IMC, LMC configurations. **C)** Example of pseudocolor maps showing the spatial distribution of granular layer responses to mossy fiber stimulation in control (*left*), 1:4 (*middle*), and 1:1 (*right*) μneedles density configurations. The triangles show the stimulating electrode location over the mossy fiber bundle (MF). GL, granular layer. PCL, Purkinje cell layer. ML, Molecular layer. **D)** The plot shows the average granular layer area responding to mossy fiber stimulation in the three conditions as in C). Data are reported as mean±mse; the superimposed *circles* indicate single data points.

**Fig. 4.**
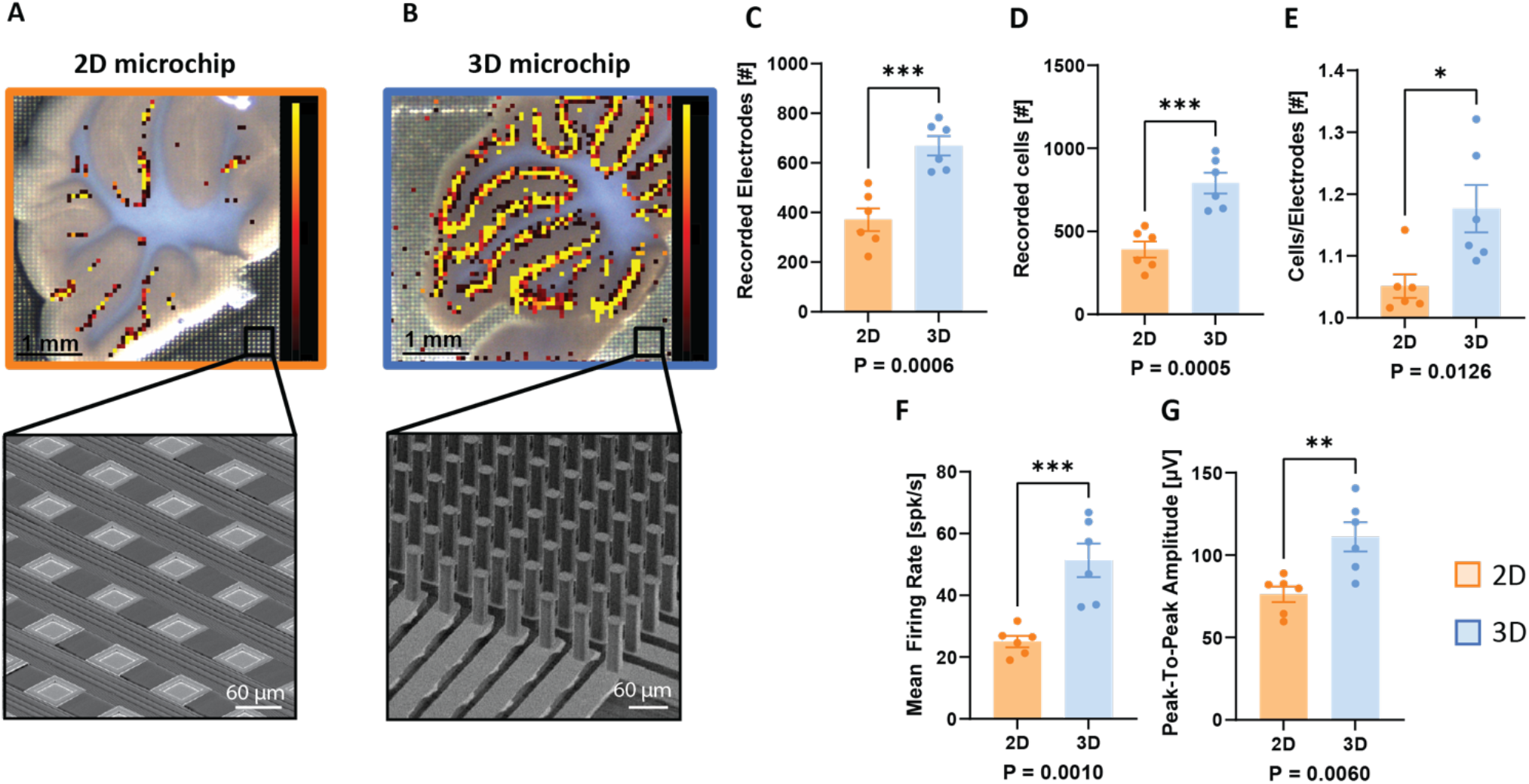
3D HD-MEA shows improved sensing capabilities when recording from cerebellar slices in comparison to planar chips. **A,B)** Optical images of slices are superimposed with activity maps showing electrodes with sustained firing rates. **A)** Cerebellar slice placed on the recording area of a planar chip reveals few active electrodes (Electrodes groups: mean firing rate >0.5 spikes/s). The magnification shows the structure of the 2D HD-MEA electrodes (scale bar: 60 μm). **B)** Cerebellar slice placed on a 3D chip reveals an appreciable activation of the entire purkinje layer across the entire slice (Electrodes groups: mean firing rate >0.5 spikes/s). The magnification shows the structure of the 3D μneedles (scale bar: 60 μm). **C – E)** The penetrating μneedles of the 3D chip allow to record from an higher number of active Electrodes (p= 0.0005, n=6), to sort a greater number of neurons (p= 0.0005, n=6) and to achieve an increase cells to electrode ratio (p= 0.0126, n=6) in comparison to the planar chip. **F,G)** The activity detected by the 3D chip is significantly higher in terms of Mean Firing Rate (p= 0.0010, n=6) and the Peak-to-Peak Amplitude (p=0.0060, n=6) compared to the planar chip. Panel A,B color scale bar: 0-10 spikes/s.

For brain spheroid experiments, spike detection was performed using the PTSD algorithm as described in Maccione et al., 2009, while spike sorting on individual electrodes was computed using a PCA with 3 features and a standard k-means clustering algorithm. A unit was considered active if the spike rate was higher than 0.1 spike/s and was located below the electrode area covered by the spheroids. In total, 6 spheroids of two different sizes were measured from a single preparation.

## Results

### *3D* μneedles *structure realization and tissue penetration*

The fabrication process described in the *Microneedle and microchannel fabrication* section has been conceived to be CMOS-wafer compatible. In this way, hundreds of chips can be structured in parallel, reducing time and cost and increasing the reproducibility of the realizations. A portion of a wafer where several chips have been structured before dicing in single units is presented in Fig. 2A. The dense structure of μneedles forms a sort of “bed of nails” where tissue remains trapped and anchored, as evident in the scanning electron microscopy (SEM) image of an entire chip in Fig. 2B. This eliminates the need to use a weight, as an omega ring, to keep the tissue in contact with the chip. A first realization with larger needles of 26 μm in width is visible in Fig. 2C. The image clearly shows as μneedles are built on larger, shorter pedestals that form a regular grid of microchannels. The close-up of the tip of one μneedle in Fig. 2D highlights the internal bulk in gold (light gray) surrounded by a thin layer of parylene. A second realization of μneedles of 14 μm in width is shown in Fig. 2E,F. Despite these structures being thinner and more flexible, they are not bent by the final diamond turning process required to open the parylene coating at the tip. Only a slight bending is in some cases visible on the μneedles at the edge of the recording area.

In terms of rigidity and penetration capabilities, the μneedles were able to access the inner layers of the tissue easily. Fig. 2G shows a SEM image of a cerebellar slice fixed on the 3D HD-MEA. The tissue was longitudinally cut using a Focused Ion Beam procedure, leaving exposed a portion of the internal layers where μneedles clearly penetrated up to a third of the total thickness of the tissue (Fig. 2H). The second set of thinner μneedles was developed for smaller and less dense biological models such as organoids or spheroids. Differential Interference Contrast (DIC) microscopy revealed that this kind of structure can enter deep into the tissue without disrupting or deforming its 3D structure. This is highlighted in Fig. 2I, where a spheroid was mounted on the 3D HD-MEA chip. The image is focused on the plane of the tips, and it is possible to observe how the tissue engulfs the μneedles without losing its spherical structure or showing evident signs of debris or cell detachment.

### Improved vitality and functional network preservation

The microchannels at the base of the μneedles allow the flowing of solution underneath the slice placed over the chip. To test whether this arrangement is able to improve tissue vitality, we performed the following tests. Cerebellar slices are an ideal model to test vitality since the high density of granule cells in the cerebellar cortex (Howarth et al., 2010; D’Angelo, 2018) makes them more susceptible to hypoxia. Acute cerebellar slices were therefore placed over chips with different microchannels width, leading to different areas in direct contact with the slice. Three conditions were tested: chips with small microchannels (SMC, with channels width of 16 μm, corresponding to a contact area on all the chip of 7.9 mm^2^); intermediate microchannels (IMC, with channels width of 24 μm, corresponding to a contact area on all the chip of 5.3 mm^2^); large microchannels (LMC, with channels width of 30 μm, corresponding to a contact area on all the chip of 3.6 mm^2^). After continuous perfusion with oxygenated Krebs solution for one hour, the slices were gently removed from the chip and loaded with 20 μM calcein AM, a non-fluorescent compound that becomes fluorescent when hydrolyzed by esterases in living cells (Braut-Boucher et al., 1995). Calcein stainings were analyzed from confocal microscopy images of each slice for both sides (Fig. 3A). The amount of living cells on the side in contact with the chip was compared to the amount of living cells on the opposite side (i.e., the side freely exposed to Krebs flowing). The decreased vitality of the side on the chip compared to the other side was then calculated. Interestingly, the decreased vitality positively correlated with the area of the slice (on the chip side) that could not access Krebs flowing (Fig. 3B). We estimated that this area is about 15mm^2^ for the control slice (no chip, directly placed over the glass of the recording chamber) and only 3.6 mm^2^ for LMC chips, with a decreased vitality of 68±4% dropping to 24±3%, respectively. These data demonstrate how the microchannels improve oxygenation and nutrient diffusion in the lower layers.

Since the tissue vitality was improved, it remained to be evaluated whether the presence of the μneedles inside the tissue might perturb network activity, for example disrupting connections and reducing the spread of signals. To address this issue, we performed voltage-sensitive dye imaging (VSDi) on cerebellar slices placed over chips with different μneedles densities. In particular, for this test, two μneedle configurations were realized: 1:1 (corresponding to 1 μneedle per pedestal, total number of μneedles 4096) and 1:4 (1 μneedle every 4 pedestals, total number of μneedles 1024). These experiments allowed us to measure the tissue area that responded to input fibers stimulation. Indeed, the cerebellar granular layer is an ideal model for detecting a possible disruption of the spatial distribution of activity. Again, granule cells are known to be densely packed, unlike other regions of the brain showing much lower cell density. Therefore, a perturbation due to the possible alteration of the three-dimensional structure of the network is more likely to be evident in the cerebellar granular layer than elsewhere. Mossy fibers provide the main input to the granular layer and are located in dense bundles in the middle of the lobules. We stimulated the mossy fiber bundle using a stimulating electrode and recorded the spread of granular layer responses using VSDi. This technique allows detecting membrane depolarization in neurons while maintaining a good level of spatial resolution (Soda et al., 2019). In this way, we calculated the percentage of granular layer area in the lobule responding to a single-pulse stimulation of the mossy fibers. We repeated this experiment on slices placed over chips with 1:1 and 1:4 μneedle densities, and without the chip (the slices were placed directly over the glass of the recording chamber, as in Soda et al., 2019). (Fig. 3C). The percentage of granular layer area responding to the stimulation was not significantly different in the slices placed over chips with 1:4 and 1:1 configurations (39.2±3.5% n=4 vs. 43.7±3.8% n=5, respectively, p=0.42). Moreover, in both cases, no difference was found compared to controls (39.7±8.1% n=5; p=0.96 and 0.66 compared to 1:4 and 1:1, respectively; Fig. 3D). These results show that the presence of the μneedles did not affect the spread of activity in the granular layer network.

Given the previous characterization, the following tests on 3D chips were performed using the 1:1 μneedle density with the LMC configuration.

### Increased sensing capabilities in detecting spiking activity on brain tissue

Once established that the 3D μneedles can efficiently penetrate the tissue without altering network properties, the signals recorded using these devices were characterized. To evaluate the sensing capabilities, 3 minutes recordings using cerebellar and cortico-hippocampal brain slices were performed, placing the same slices on a 2D planar chip and then transferring them to the 3D chip (or vice versa for half of the slices used). Fig. 4A,B shows an illustrative example of a cerebellar slice positioned firstly on a planar HD-MEA (a) and then on the 3D HD-MEA (b). From a qualitative analysis of the firing rate maps, it was evident the improved sensitivity of the 3D HD-MEA in detecting activity from the Purkinje cell layer compared to a planar HD-MEA. The activity map, indeed, perfectly matched the anatomical structure of the tissue highlighting the lobule positions of the entire brain slice.

From a quantitative perspective, in cerebellar slices, the electrodes having a mean firing rate higher than 0.5 Hz were considered to represent the active area of the slice. The recordings performed on the 3D chip showed a significant increase in the number of active electrodes (p = 0.0006), the number of sorted cells (p = 0.0005), and the cells to electrodes ratio (p = 0.0126), compared to the same slices on the planar chip (Fig. 4C,D,E). Further investigations were performed to understand if the penetrating μneedles were significantly more effective in detecting the spiking activity of the cerebellar slices. After spike detection and sorting analysis, a significantly higher mean firing rate in the cerebellar slices placed in the 3D chip was observed compared to the 2D (p = 0.0010), together with a significant improvement of the peak-to-peak amplitude (p = 0.0060) (Fig.4F,G).

In cortico-hippocampal slices, a 100Hz high-pass filter was applied to remove slow oscillations and record only the spiking activity. Only the electrodes having a mean firing rate higher than 0.1 Hz, restricted only to the cortical region, were considered for the analysis (Fig. 5A,B). Similar consideration in terms of improved activity in the spike maps can be extrapolated as for Fig. 4A,B. The penetrating μneedles of the 3D chips allowed to record a significantly higher number of active electrodes in the cortical area with respect to the planar chip (p = 0.0384) as shown in Fig. 5E. Consequently, this resulted in a significantly larger number of spikes detected in the slices placed on the 3D chip (p = 0.0348) (Fig. 5C,D). However, the peak-to-peak amplitude did not change between the waveforms detected by the 3D and the planar HD-MEAs (Fig. 5F). Considering that cortical activity is not autorhythmic and is characterized by a more “random” and sparse activity compared to Purkinje cells, the data suggest that the 3D chips couple better the tissue and access to inner areas where neurons are better preserved by possible damages induced by the cut, thus being able to reach a larger number of active neurons.

**Fig. 5.**
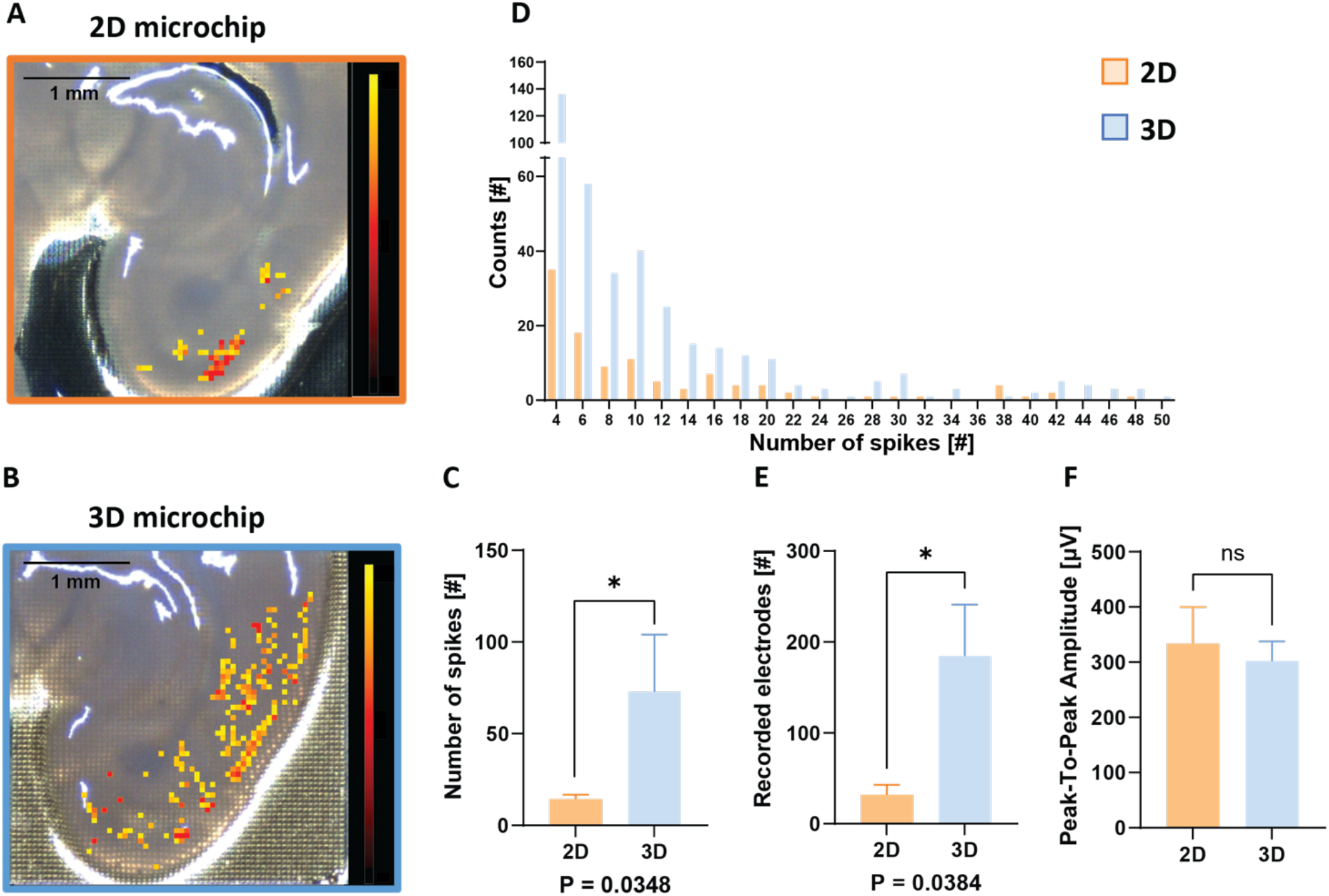
3D HD-MEA improves sensing capabilities also in a cortico-hippocampal slice model with respect to a planar HD-MEA. **A,B)** Cortical area exhibits rare active electrodes when recorded with a planar chip (A) with respect to the 3D which shows a more spread and uniform activation on the entire cortical region (B). For both panels, electrodes groups: mean firing rate >0.1 spikes/s. **C,D)** The number of spikes detected by the perforating μneedles is significantly higher than the 2D chip (p= 0.0348, n=4). **E,F)** Despite the higher number of recorded electrodes obtained with the 3D chip (p= 0.0348, n=4) as shown in E, no sensing improvement in terms of Peak-to-Peak amplitude was observed (F). Panel A,B color scale bar: 0-10 spikes/s.

In aggregate, these data show that the μneedles of the 3D microchip significantly improve the sensing capabilities of the electrodes, regardless of the specific area involved.

### Accelerating compound effect in modulating electrical activity

The presence of a grid of microchannels at the bottom of the 3D HD-MEA chip was already demonstrated to increase cell viability and tissue integrity due to the possibility of directly oxygenating the bottom cell layers. The microchannels might also improve compound diffusion, reducing the time of action of a substance in modulating electrical activity. To prove that, spontaneous activity of cerebellar slices was recorded for one minute before perfusing Krebs with 3μM TTX. Fig. 6A,B show a qualitative example of the complete abolition of the spiking activity after 2.5 min TTX application as recorded by a 3D HD-MEA. Both in slices placed over a 2D and a 3D HD-MEA, the decay was evident and quite sudden. Nevertheless, slices on 3D chips showed a faster decrease of the mean firing rate after TTX application, reaching an overall depression of neuronal activity significantly earlier than slices on planar chips (Fig. 6C). In particular, cerebellum slices recorded on the microchannels grid needed less time to reach 10%, 50%, and 90% decreased activity with respect to the baseline as shown in Fig. 6D. These results demonstrate a clear improvement in the efficiency of the compound action using the 3D chip.

**Fig. 6.**
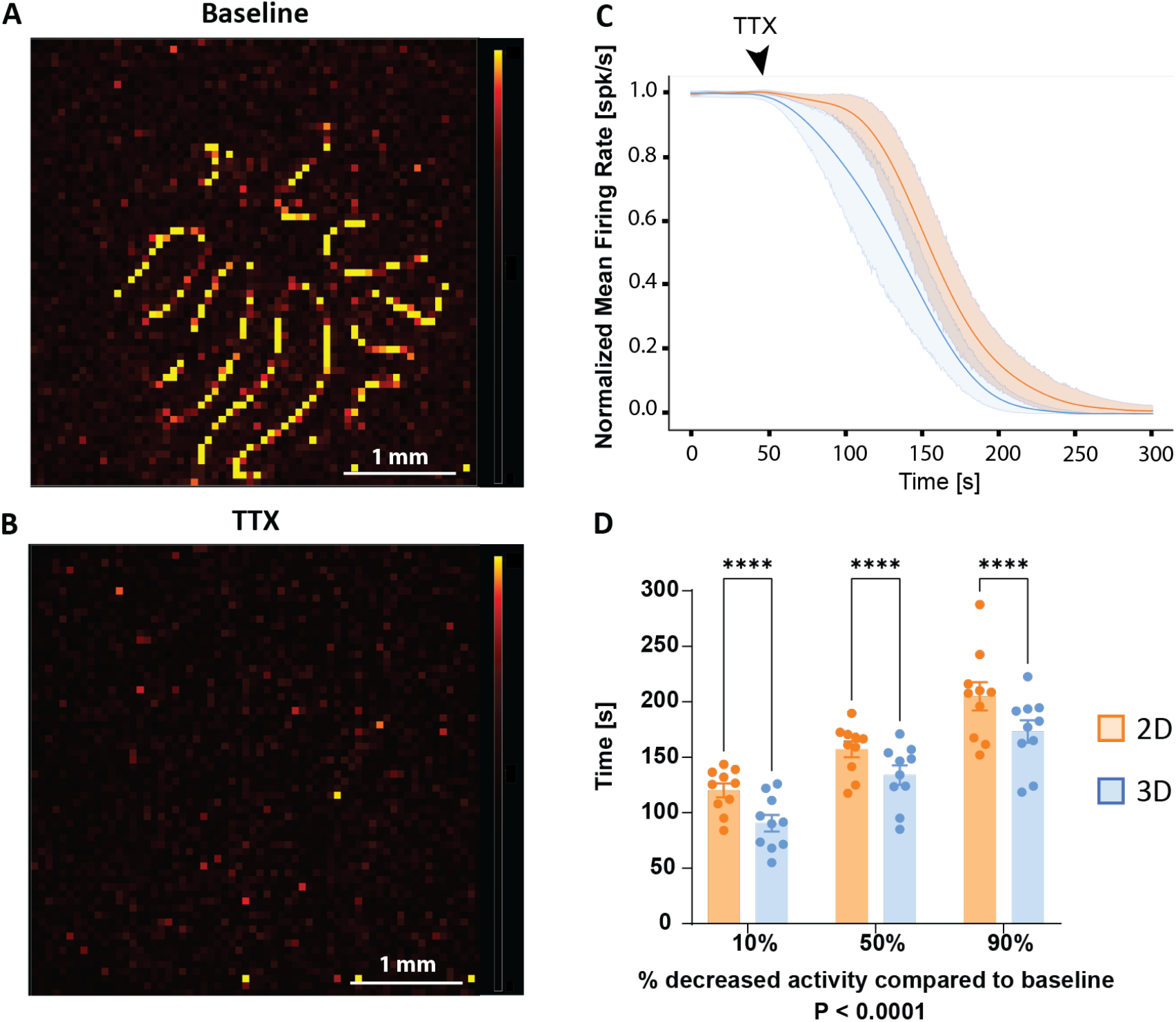
Improvement in the efficiency of TTX modulation on cerebellar slices recorded with 3D chips. **A,B)** The activity maps show the active electrodes before (A) and after the TTX application (B): the peculiar lamellar shape of the cerebellum slice completely disappears consequently to TTX perfusion indicating a massive depression of the activity. **C)** Cerebellar slices placed on 2D and 3D chips display the same decay trend of the neuronal firing due to TTX treatment, but with an anticipated and faster decrease in the activity observed using the 3D technology. Activity was normalized with 1 min basal activity before compound application. **D)** Time to reach 10%, 50%, 90% of decreased activity for 2D and 3D HD-MEA after TTX application. The firing activity requires significantly less time to decrease 10%, 50%, and 90% when tested on the 3D HD-MEA chips which integrate a grid of microchannels to improve compound diffusion (10%, %50%, 90%, p<0.0001, n=10 for both conditions). Panel A,B color scale bar: 0-300 μV.

### Stimulating capabilities

Both the planar and the 3D chip have the ability to release an electrical stimulation from every single electrode, allowing the detection and the recording of evoked activity from many experimental models. However, in cerebellar slices, the response to the stimulus is particularly fast and challenging to detect. Furthermore, efficient stimulation often requires penetrating the tissue to access inner mossy fibers. For these reasons, the planar chip is often inefficient in inducing and identifying such evoked activity. Contrarily, the μneedles that characterize the 3D technology proved to be efficient in reaching the mossy fiber bundle and detecting the granular layer response with exceptionally appreciable timing and spatial resolution. To evoke neuronal responses, a biphasic stimulus was released in different cerebellar lobules of the same slice (see *Electrophysiological recordings* Section). An evoked signal propagation has been identified in each stimulated lobule, as shown in Fig. 7A, with an average calculated speed of 311 ± 39 mm/s (n= 8). Robustness of the response in terms of signal amplitude and duration can be appreciated in the exemplificative raw traces in Fig. 7B. Notably, with planar HD-MEAs it was not possible to obtain similar results, thus demonstrating the higher enhanced stimulating capabilities of the penetrating μneedles compared to flat electrodes.

**Fig. 7.**
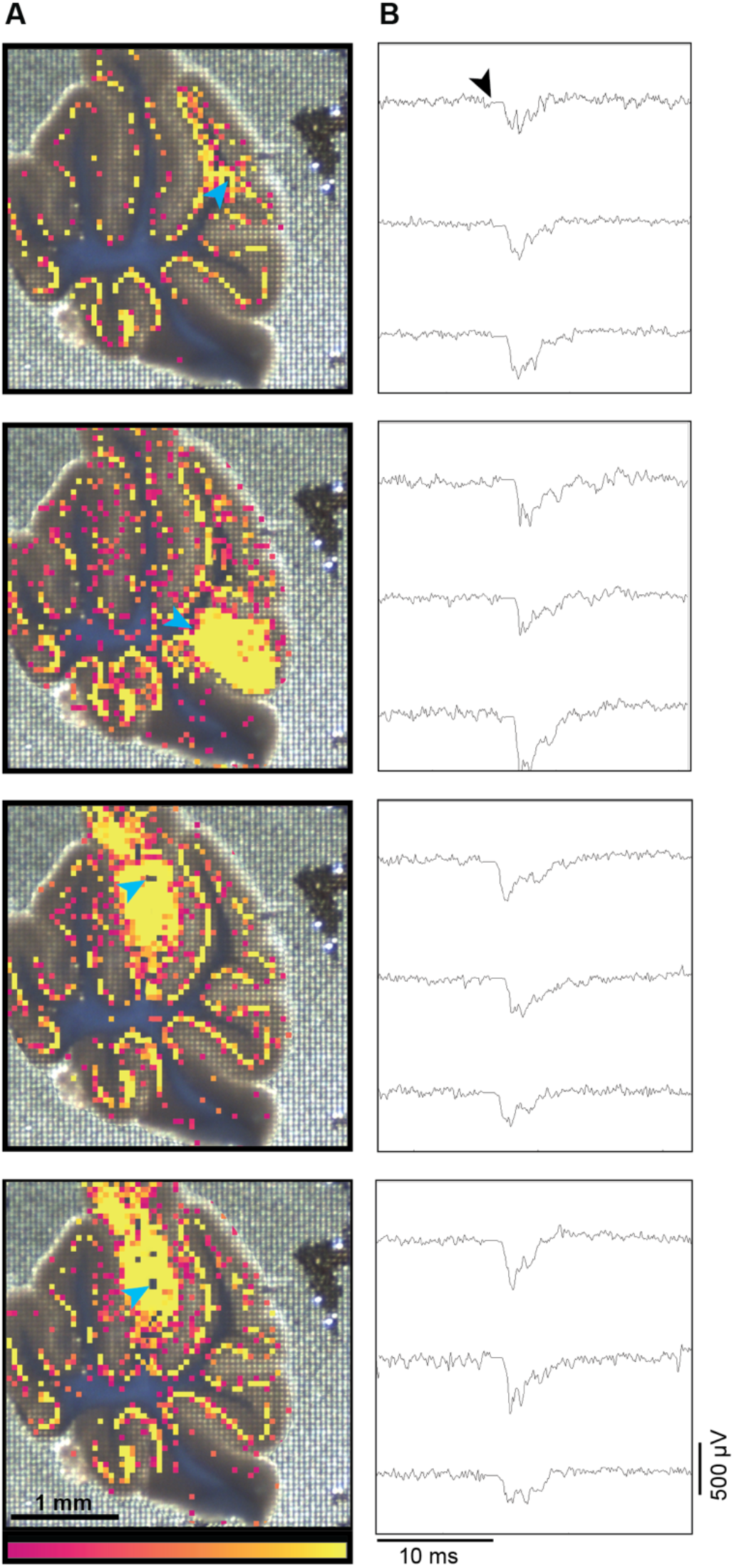
Robust evoked activity resulting from the electrical stimulation of a cerebellar slice performed with the 3D technology. **A)** Stimulation of the mossy fiber bundle in different lobules, at the specific locations indicated by the blue arrows, on the same cerebellar slice. Area activated upon stimulation matches the anatomical organization of the tissue. **B)** Examples of single raw traces from three different electrodes per lobules, showing the local field potentials recorded in the granular layer in response to mossy fiber stimulation. Notice that, since granule cells are silent at rest, only the granular layer of the stimulated lobules is active. Purkinje cells are evident in all the lobules due to their autorhythmic activity. Panel A color scale bar: 0-10 spikes/s.

### Measuring activity from spheroids

The need to generate *in vitro* cell cultures capable of maintaining a 3D structure is a challenge that, in recent years, has led to the creation of increasingly complex models (Kelava and Lancaster, 2016). However, measuring network activity in such models is still problematic, mainly due to the difficulty in accessing the internal boundary of the cell model. A second set of thinner μneedles was fabricated to penetrate low density 3D structures without damaging them. This set was tested with brain spheroids from primary embryonic neurons, cultured in two different sizes of about 400 μm and 600 μm in diameter. The large recording area (3.8 by 3.8 mm^2^) allowed the positioning of multiple samples on the electrode array, and, at the same time, the high spatial resolution confers the advantage of maintaining a high number of recording sites. For these reasons, three spheroids sorted by size were mounted on two different chips. Acute measurements obtained immediately after placing the spheroids revealed the presence of robust spiking and bursting activity (Fig. 8A,B right panels). The left panels in Fig. 8A,B shows the three spheroids in the center of the recording area and the active electrodes beneath. Despite the fact that the three adjacent spheroids were sharing the same media, their activity was not synchronized, as evident in the raster plots in Fig. 8A,B (middle panels). This observation indicates that each element can be considered, during the analyses, as a single, independent sample.

**Fig. 8.**
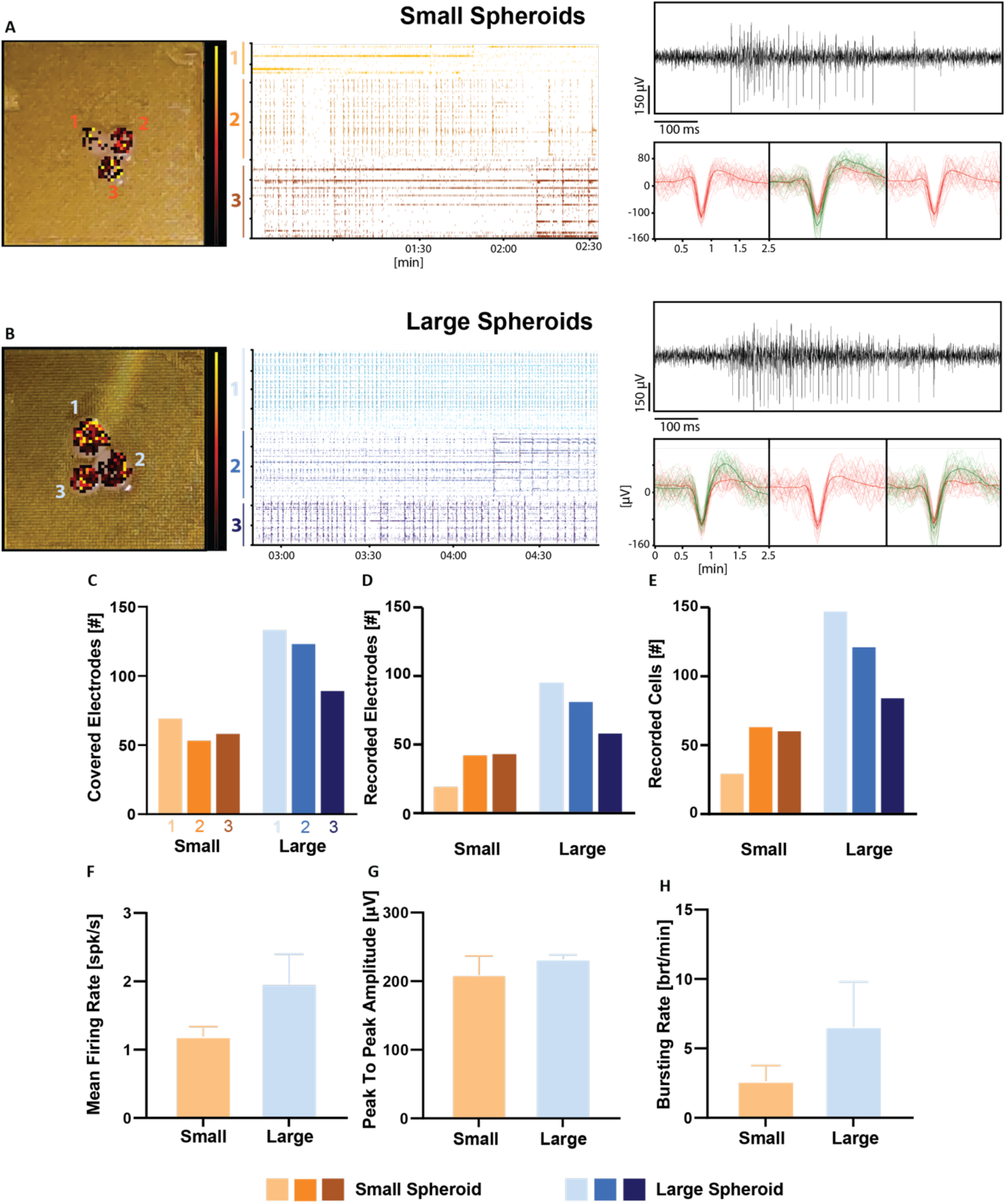
3D HD-MEA equipped with a small set of μneedles (width 14 μm, height 65 μm) are optimized for spheroid activity measurements. **A,B)** Picture of the 3 spheroids on chip with active channels (*left*), raster plot of the spiking activity over 2.5 minutes (*middle)* and representative images of a raw trace and averaged waveforms for three electrodes (*right)*. **C-E)** The plots show: **C)** the number of electrodes covered by the spheroids, obtained superimposing the picture on the electrode array; **D)** the number of recorded electrodes (i.e., all the electrodes containing at least 1 active cell after spike sorting); **E)** number of recorded/active cells (i.e., with mean firing rate ≥ 0.1 spikes/s). **F-H)** Differences in the activity between small and large spheroids were observed by measuring the mean firing rate **F)** and the bursting rate **H)**. The size of the spheroids seemed not to affect the peak-to-peak amplitude **G)**. Panel A,B color scale bar: 0-10 spikes/s.

Large spheroids were penetrated by more μneedles (Fig. 8C) and, consequently, they showed a larger number of electrodes (Fig. 8D) where active cells were detected (Fig. 8E). However, the ratio “active cells/covered electrodes” was similar for both large and small spheroids (<1.5, data not shown), thus demonstrating that the size of the models did not directly affect the sensing/penetrating capability of the μneedles. The results extracted from every single spheroid were averaged to create experimental groups divided by spheroids sizes. The mean firing rate, bursting rate, and peak-to-peak amplitude of the two groups are shown In Fig. 8F-H. It is interesting to notice that, unlike for peak-to-peak amplitude (Fig. 8H), larger spheroids show a trend towards an increased mean firing rate and bursting activity (Fig. 8F,G), though not statistically significant. This suggests that the size of the spheroids does not impact either the signal amplitude or the neuron-electrode coupling.

## Discussion

The use of MEAs on 3D models is constantly growing, thanks to the possibility of recording extracellular electrical activity from a network of cellular elements, as in slices and organoids. Nevertheless, some relevant issues of technical and biological origins limited the potentiality of this approach. First, the side of the tissue in contact with the recording probes suffers the absence of direct perfusion of the medium in the recording chamber. Nutrients and gasses need to diffuse through the thickness of the tissue to allow an exchange between cells and the external medium. This is also true for any molecule added to the bulk perfusion, and their effects might be reduced or slower than optimal. Moreover, the outermost areas of a 3D model often show dead cell layers due to methodological issues, as evident in the slicing procedure, where this phenomenon can be reduced but not solved using advanced vibroslicers. Herein, the implementation of a 3D HD-MEA chip capable of overcoming all these issues and providing additional crucial advantages was presented in this paper.

### Fabrication process

Despite the fabrication process of the 3D HD-MEA relying on well-established clean room techniques, the entire chain requires a large number of steps, and it has been designed to be compatible with the CMOS wafer. Therefore, each procedure was developed in order not to stress the microcircuitry below or to damage the contact pads of the chips. Furthermore, the fabrication process was developed to structure an entire wafer, thus producing hundreds of pieces at the same time. This has an evident impact in terms of reliability and reproducibility of the chip, making this implementation potentially scalable and giving the possibility to make the 3D HD-MEA technology accessible by any lab.

### Overcoming the main MEAs-related issues

Data demonstrate that the 3D μneedles penetrate the tissue, gaining access to the inner structures beyond the dead cell layer. This feature was tested on cerebellar slices, where the high cell density is ideal for this purpose, and obtained optimal results, as evident in Fig. 2H. At the same time, the presence of a microfluidic system at the base of the chip significantly improved slice vitality, as shown by calcein assays. Interestingly, the vitality preservation correlated well with the microfluidic system area. The efficiency of this arrangement is evident when testing drug perfusion in the slice, showing a marked decrease in the effect kinetics on 3D vs. 2D chips. At this point, the main concern might be the possible perturbation of the tissue integrity once placed over the μneedles. Therefore, network activity in the granular layer was tested using VSDi on cerebellar slices. Here, given the high cell density of this layer, μneedles can potentially cause more damage to network integrity than in other brain areas. Our data showed that, upon stimulation of the mossy fiber bundle, the spread of the network response resulted in an activated area that is not altered by μneedles presence in the tissue, both at full (1:1) and reduced (1:4) density. Taken together, this first set of data demonstrates that the use of these chips determines increased tissue vitality and provides access to the tissue beyond the dead cell layers with efficiently penetrating μneedles without affecting neuronal network structure and function. As a result, the testing of 3D chip activity was performed on chips with the 1:1 configuration with larger microchannels.

### Providing crucial advantages

Besides improving tissue vitality, the 3D chip architecture provides several crucial advantages. First, the close relationship between the recording probes and the cellular source of the signal significantly increases the efficiency of signal acquisition, which reflects on the number of active electrodes, the cells to electrodes ratio, and the number of recorded units. Secondly, the signal-to-noise ratio is significantly improved in this configuration. Thirdly, each of the 4096 μneedles can be recruited for the stimulation of the sample. Given the high μneedles density, the stimulation site can be determined with a precision of a few tens of μm, and the stimulated area can be customized with very few constraints. As expected, the electrical stimulation of a 3D tissue by penetrating μneedles is far more efficient than that provided by planar probes. In addition, the improved sensing capability allows the detection of small and fast responses, such as the cerebellar ones. Even more notably, this system can stimulate and record network responses in different areas of the slice, exploiting the high spatial definition. In the example in Fig. 7, different cerebellar lobules were stimulated, making it possible to characterize their responses in the same slice. This offers the unique advantage of comparing network and subnetwork properties in the same brain area, in the same slice, discerning the properties of separate modules or microcircuits. Moreover, by increasing the efficiency of molecule diffusion and improving their action kinetics, this new 3D chip is the ideal tool to perform fast and efficient drug testing and screening on new 3D models such as organoids and neural spheroids, where recordings from multiple samples are possible at the same time (see Fig. 8A,B). Finally, the 3D chip does not require any anchor or platinum ring to keep the sample in place during the recordings. The μneedles themselves stabilize the tissue, improving the reliability of recordings, especially in the long term. Indeed, this configuration abolishes any anchor-related tissue stretch, avoiding possible damages and artifacts to cell functioning. As a final remark, it is worth mentioning that increasing the efficiency of the technical apparatus will determine, as a consequence, a reduction in the number of tests (and therefore animals) needed for obtaining statistically relevant results. Such technological advancements are crucial in realizing the 3Rs principles.

## Conclusions

The technological advancement characterized in this paper has the potential to be a game changer in neurophysiological research, besides other fundamental applications on excitable tissues (e.g., muscle and heart physiology, non-brain organoids). It provides an unprecedented combination of state-of-the-art temporal and spatial resolution of extracellular recordings, powerfully combining increased vitality of the sample and tissue penetration, optimizing the signal-to-noise ratio and the efficacy of recordings in terms of cellular units at the source of recorded signals. The perspectives opened by such a technology are quite broad: from drug testing on living tissues to high precision network activity reconstruction, using advanced statistical methods to determine circuit connectivity and complex dynamics. Furthermore, the amount of information derived from single recordings is unprecedented and opens the door to computational applications. Interestingly, theoretical models can be helpful in further interpreting the data (Schepper et al., 2021). At the same time, the massive amount of data from simultaneous recordings of thousands of neurons in the same sample is ideal for instructing computational models of neuronal networks. This application is particularly intriguing given the recent boost in this field, which needs experimental evidence for reconstruction and validation.

## Author Contributions

ED, LM, ST, AMa, MT, MG conceived the experiments

LM, AMa, MT, GS, CB, ST, AMo, AO performed the experiments

OD, FC, PS, JS, KI produced and electrically tested the 3D HD-MEA

LM, AMa, GS, CB, ST analyzed the data

CC, MM provided experimental support

ED, AMa, LM, GS, MT, CB wrote the paper

All the authors revised the paper

## Acknowledgments

The authors want to thank Dr. Michele Dipalo at the Italian Institute of Technology (IIT, Genova Italy) in providing SEM images of a cerebellum slice on a 3D HD-MEA chip shown in Fig. 2.

